# Unbiased identification of nanoparticle cell uptake mechanism via a genome-wide CRISPR/Cas9 knockout screen

**DOI:** 10.1101/2020.10.08.332510

**Authors:** Eric L. Van Nostrand, Sarah A. Barnhill, Alexander A. Shishkin, David A. Nelles, Eric Byeon, Thai Nguyen, Yiu Chueng Eric Wong, Nathan C. Gianneschi, Gene W. Yeo

## Abstract

A major bottleneck in nanocarrier and macromolecule development for therapeutic delivery is our limited understanding of the processes involved in their uptake into target cells. This includes their active interactions with membrane transporters that co-ordinate cellular uptake and processing. Current strategies to elucidate the mechanism of uptake, such as painstaking manipulation of individual effectors with pharmacological inhibitors or specific genetic knockdowns, are limited in scope and biased towards previously studied pathways or the intuition of the investigators. Furthermore, each of these approaches present significant off-target effects, clouding the outcomes. We set out to develop and examine an unbiased whole-genome screening approach using pooled CRISPR/Cas9 libraries for its ability to provide a robust and rapid approach to identify novel effectors of material uptake. Enabling this, we developed a methodology termed fast-library of inserts (FLI)-seq for library preparation and quantitative readout of pooled screens that shows improved technical reproducibility and is easier to perform than existing methods. In this proof-of-concept study we use FLI-seq to identify a solute carrier protein family member, SLC18B1, as a transporter for polymeric micellar nanoparticles, confirming the viability for this approach to yield novel insights into uptake mechanisms.

## Introduction

Methods for intracellular transport of biomolecules are much sought after in the context of both *in vitro* delivery reagents and *in vivo* therapeutics (1–4). Recently, we found that micellar assemblies of hundreds of amphiphiles consisting of single-stranded DNA which has been covalently linked to a hydrophobic polymer, referred to as DNA-polymer amphiphile nanoparticles or DPANPs, can readily access the cytosol of cells where they modulate mRNA expression of target genomes without transfection or other helper reagents, making them potential therapeutic nucleic acid carriers (5, 6). However, despite their effective uptake properties and efficacy in the cytosol, it was unknown how these polyanionic structures can enter cells. Indeed, generally, bottlenecks in understanding and achieving delivery and uptake remain a forefront issue in translatability of macromolecular and nanomaterials-based therapeutics generally, including with respect to nucleic acid therapies (7). However, elucidating these mechanisms is typically done through a collection of poor, low throughput choices that include stepwise knockouts of genes postulated to be potential candidates (e.g. anion scavengers) (8), the use of pharmacological inhibitors that often have broad side-effects, or membrane disruption methods that raise questions as to biological relevance (9). Thus, unbiased screening technologies for the identification of molecular regulators of uptake would dramatically improve our ability to develop and understand synthetic molecule and material uptake and transport (10, 11). Advances here, capable of human genome-wide analysis, would represent a breakthrough in probe and therapeutic development by unlocking the capability to rationally design structures that interact with and internalize into cells via specific, targeted mechanisms.

Forward genetics has long been a powerful approach in the biologist’s toolbox to identify functional regulators of undeciphered mechanisms. Specifically, the development of pooled shRNA and more recent CRISPR/Cas9 screens have provided a strategy to characterize thousands of specifically targeted manipulations in a single experiment (12). Standard CRISPR/Cas9 targeting involves expression of both Cas9 protein and a single guide RNA (sgRNA) which directs double-strand cleavage of a targeted DNA region, leading to insertions or deletions (indels) in the target region (13). Pooled screening is frequently performed with a CRISPR library containing single guide (sg)RNAs targeting hundreds to tens of thousands of annotated genes, typically with multiple guides per gene, to generate a population of cells representing individual knockout clones (14–16). Cells are then subjected to a selective pressure or treatment (such as treatment with Cy5- or other fluorophore-labeled DPANPs), and genomic DNA isolation and library preparation followed by high-throughput sequencing quantifies the changes in sgRNA frequency between selected sub-populations to identify gene knockouts that show altered phenotypes between distinct conditions, timepoints, or treatments as desired. However, as part of performing these screens, we found that amplifying the library without introducing bias into the relative target frequencies was a complex challenge with numerous sources of technical variability during this process ranging from amplification biases due to GC-content and primer positioning, and secondary structure (17, 18). In particular, the amount of genomic DNA itself can be a significant limitation, as 100-fold coverage of a standard 60,000 sgRNA pool requires amplifying approximately 40 μg of genomic DNA, but amplification inhibition is often seen in PCR with more than 500 ng – 2 μg per reaction. A variety of methods have been described to alleviate this concern, including simply performing multiple (often 30 or more) PCR reactions per sample, performing restriction digestion and size-selection to remove unrelated genomic DNA (19), and pulldown with locked nucleic acid probes to specifically enrich for the region of interest (20). However, these approaches scale poorly with high sample amounts, limiting the ability to perform replicates or explore additional sub-populations.

Here, we describe our development of a simple, low-cost approach for targeted amplification of CRISPR/Cas9 and other pooled screens using biotin-coupled RNA oligonucleotides that shows near-complete recovery of desired sgRNA target regions while removing unwanted genomic DNA background, allowing low-amplification library preparation at scale and with decreased technical variability. The development of this approach enabled us to perform a genome-scale CRISPR/Cas9 screen to discover modulators of DPANP uptake in HEK293T cells. Confirming the value of this unbiased approach to the study of material transport, we identify a solute carrier protein transporter (SLC18B1) as a novel effector of DPANP uptake. Thus, the use of pooled screening approaches can provide novel insights into as yet uncharacterized mechanisms of transport of therapeutically relevant materials and molecules, creating unique opportunities for future therapeutic development.

## Results

### Simplified library preparation for readout of pooled screens

We set out to use genome-wide CRISPR/Cas9 knockout screens to identify novel regulators of DPANP uptake in HEK293T cells. However, initial experiments showed poor reproducibility across biological and technical replicates, matching previous observations in re-analysis of published CRISPR screens (21). Thus, there is a need for improved experimental approaches in accurately and quantitatively assaying significant effectors.

To develop an improved method for readout of pooled CRISPR/Cas9 screens, we performed a genome-wide CRISPR/Cas9 knockout screen in 293T cells using the published GECKO v2 ‘A’ and ‘B’ libraries, which contains 3 guides targeting each of 19,050 genes for a total of 63,950 and 56,869 unique guides respectively (including 1,000 non-targeting controls (NTCs)) (Fig. 1a) (16). For each replicate, 36 million 293T cells per experiment were transduced with lentivirus at 0.3 multiplicity of infection, after which cells were grown under puromycin selection for 10 days with at least 100x coverage to ensure infection and high knockout efficiency. Genomic DNA (gDNA) was isolated at D0 (immediately after lentiviral spinfection), and 3, 6, and 10 days after infection to assay dropout of essential genes, with an average of ~9.4 μg gDNA recovered per million cells, matching standard expectations for aneuploid HEK293T cells (Sup. Fig. 1a). Thus, to maintain 100x coverage of the sgRNA library would require library preparations with ~60 μg gDNA. As previous described, using the standard method of simple PCR amplification with targeted primers (14) we observed inhibition of PCR at ~2 μg of gDNA per PCR (with a 2X master mix formulation showing the least inhibition, but the limited volume available per reaction limits the maximum gDNA in this case) (Sup. Fig. 1b). Although this is consistent with standard inhibitory effects of excess DNA, it suggests that proper amplification would require dozens or more separate amplification reactions per sample.

**Figure 1.**
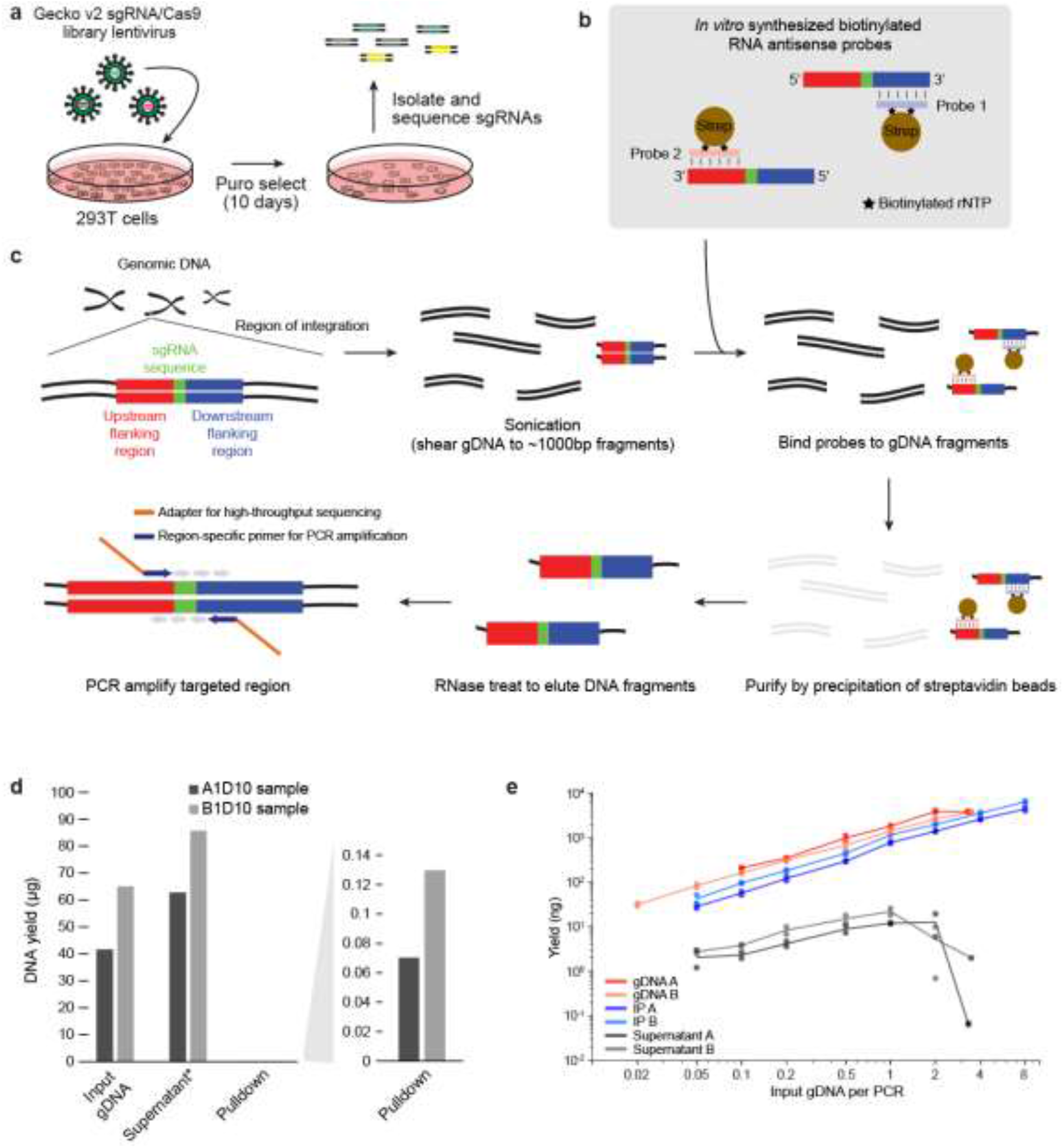
Improved library preparation for CRISPR/Cas9 pooled screens. (a) In pooled CRISPR/Cas9 screening, first a pool of lentivirus is made in which each lentiviral particle contains sgRNAs targeting one of hundreds to thousands of genes. After infection at low multiplicity of infection to ensure at most one lentiviral particle infects each cell, cells are selected for viral infection for 10 days. In a simple screen for lethality, gDNA is then isolated and queried to look for sgRNAs which are no longer present (indicating that the targeted gene is essential). (b) RNA antisense probes were generated from PCR amplification of ~500nt fragments of constant regions flanking the variable sgRNA region, followed by T7 *in vitro* transcription with biotinylated rUTP or rCTP. (c) To deplete non-sgRNA containing regions, genomic DNA is isolated and sonicated. Antisense probes are then bound to gDNA fragments, followed by purification with streptavidin beads. RNase treatment is then used to remove RNA probes and isolate ssDNA fragments containing sgDNA regions, followed by PCR to add adapters for high-throughput sequencing. (d) Bars indicate enrichment performed on biological replicate Day 10 samples (A = ~4.4×10^6^ cells, B = ~6.9×10^6^ cells). Shown are input gDNA (dsDNA), supernatant (ssDNA), and pulldown (ssDNA) yields quantified by Nanodrop 2000. (e) Points indicate (y-axis) yield (quantified by high-sensitivity D1000 tapestation) after 20 cycles of PCR amplification starting with (x-axis) indicated input gDNA amounts (or equivalent sample fraction for purified and supernatant samples).

To address this challenge, we developed the Fast Library of Inserts (FLI-seq) approach to remove the majority of unwanted non-sgRNA genomic DNA (Fig. 1b-c). Multiple such approaches exist, with restriction digest followed by DNA gel electrophoresis and size selection the most commonly used approach (19). However, due to the handling complexity and inefficiency of this approach at large scales, we tested an alternative approach in which we performed pulldown with biotin-labeled antisense RNA oligonucleotides to selectively enrich sgRNA-containing regions relative to overall gDNA. We generated antisense probes by PCR amplification of a 500nt constant region flanking the sgRNA sequence, followed by *in vitro* T7 transcription with biotinylated UTP (Fig. 1b) (see Methods). After incubation, streptavidin pulldown, and washes, we found that 99.8% of input gDNA remained in the supernatant, with less than 0.2% purified after enrichment (Fig. 1d). However, when PCR was performed with DNA amounts equivalent to equal numbers of input cells, we observed that the enriched fraction gave similar yields to non-selected gDNA and far greater than supernatant (Fig. 1e). Using values from 0.5 μg-equivalent input material, this corresponds to a ~30-fold increase in amplified yield between enriched and supernatant, indicating a greater than 10^4^-fold enrichment for sgRNA regions in pulldown material (Fig. 1e). Furthermore, we were able to increase input material to the equivalent of 8 μg total gDNA (=8.5×10^5^) cells per 20 μL PCR reaction without observing noticeable drops in yield (Fig. 1e), indicating that this approach can yield robust amplification with far fewer PCR reactions necessary. We note that the enrichment procedure includes an additional column cleanup step, which may explain a large fraction of the ~2.5-fold decreased yield relative to non-selected gDNA; similar loss would be expected from post-PCR restriction digest and gel electrophoresis steps utilized in other protocols.

**Supplemental Figure 1.**
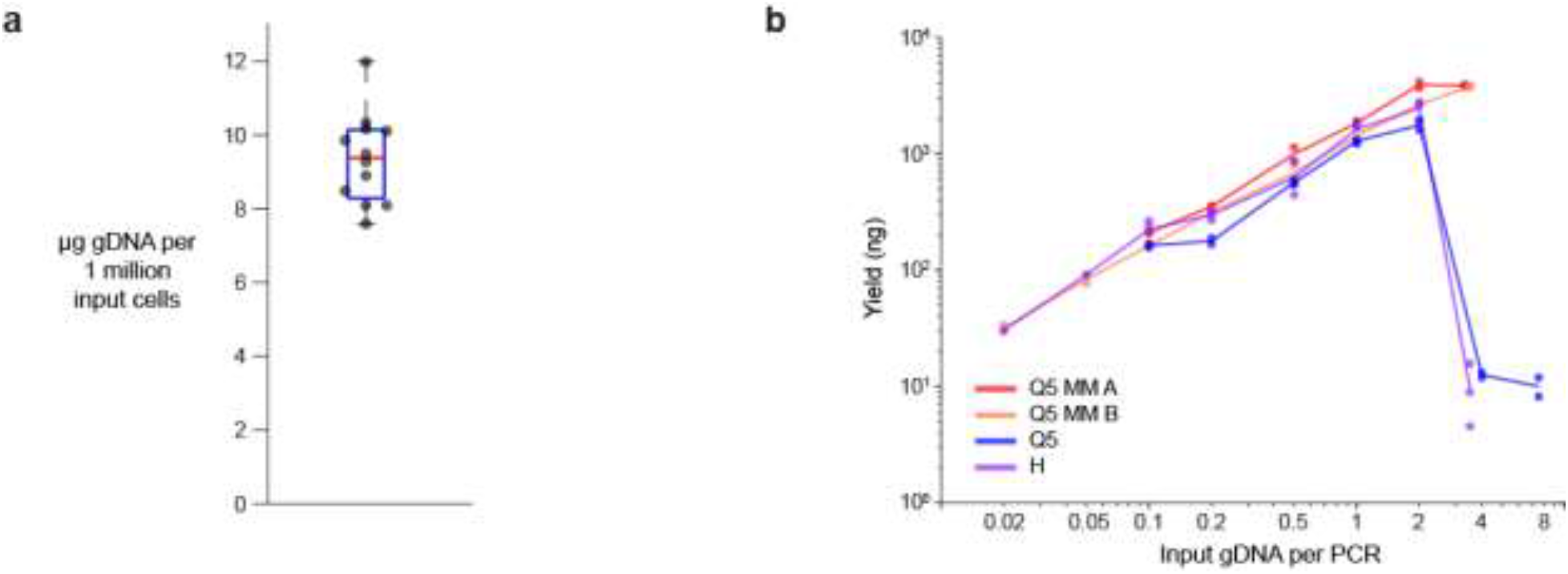
Improved library preparation for CRISPR/Cas9 pooled screens. (a) Points indicate gDNA yield per million HEK293T cells from 12 biological replicate experiments. Boxplot indicates 25^th^ to 75^th^ percentile, with median indicated in red and whiskers extending to outlier points. (b) Points indicate (y-axis) DNA yield after 20 PCR cycles of amplification for (x-axis) various gDNA amounts. Shown are two biological replicates (red) using Q5 2X master mix, (blue) Q5 Hot Start, and (purple) Herculase II Fusion DNA Polymerase PCR reagents.

### Validation of simplified library preparation reproducibility

Although this approach significantly decreases the necessary number of PCR reactions and handling complexity, it is critical that the approach properly enriches for sgRNAs that are enriched in the cellular pool and does not introduce high levels of technical irreproducibility. First, to query whether we recover true signal, we performed library preparation and high-throughput sequencing of gDNA from D0 and D10 post-infection of HEK293T cells. We observed that guides targeting essential genes (22) showed significant depletion by D10, confirming successful integration and excision by Cas9 and successful library generation (Sup. Fig. 2a).

Next, to assay reproducibility, we defined two replicate structures: infection replicates (in which two replicate infections were performed and cells were maintained separately through the entire experiment) and technical replicates (in which after 10 days in culture, isolated genomic DNA was split in half and independently enriched and amplified into library) (Fig. 2a). After processing reads to quantify normalized sgRNA read density, we observed high concordance for technical replicates at 100x coverage (R = 0.94 between log_2_(RPM)), confirming that the FLI-seq approach reproducibly recovers sgRNAs with low variability (Fig. 2b-c, Supplementary Table 1). Further, we observed that infection replicates showed similarly high concordance (R = 0.87), indicating that much of the variability commonly attributed to infection may simply be due to technical library preparation variation (Fig. 2d). Next, we queried the effect of beginning enrichment with variable library coverage by varying the input gDNA amount from 12x to 200x coverage (Fig. 2a). We observed correlations of 0.88 or above between technical replicates for all samples with at least 50x coverage (~3 million cell input), with 50x technical replicates showing less error than 100x infection replicates (R = 0.89 vs 0.87 respectively) (Fig. 2e). However, significant variability observed when gDNA input was less than 35x, suggesting that bottlenecking remains a significant concern at this low coverage level (Fig. 2f).

**Figure 2.**
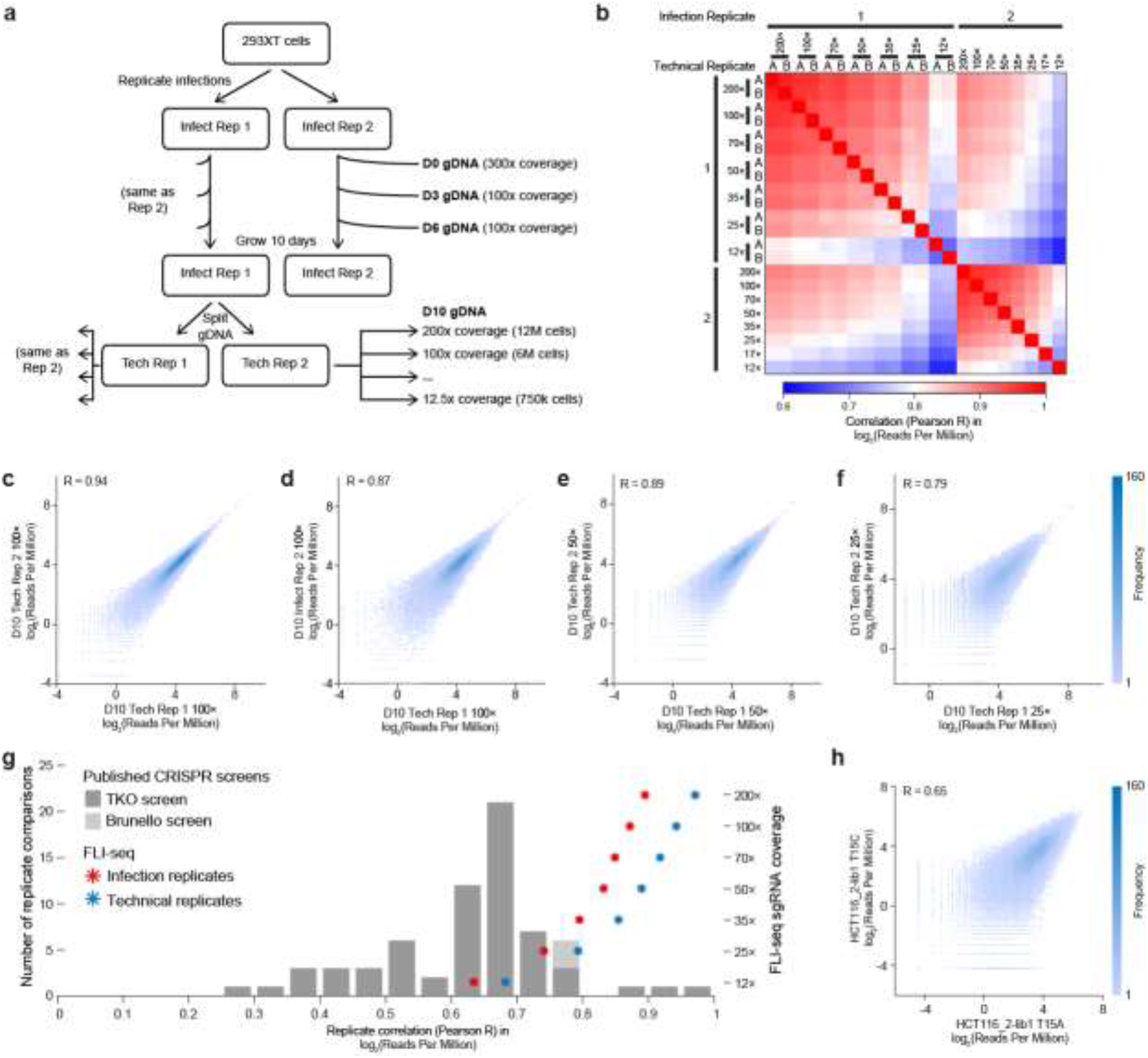
Reproducibility testing for improved library preparation for CRISPR/Cas9 pooled screens. (a) Schematic of infection and technical replicates to query experimental reproducibility. Coverage was extrapolated based on gDNA yield using standard 6.6 *μ*g per 1 million cell estimates. (b) Heatmap indicates correlation (Pearson R) between reads-per-million normalized sgRNA counts (log_2_) from technical or biological replicate samples as indicated, using only sgRNAs with RPM>1 in at least one dataset. (c-f) Scatter plots indicate frequency of sgRNAs with indicated read coverage for (c) D10 100X coverage technical replicates, (d) D10 100X coverage infection replicates, (e) D10 50X coverage technical replicates, and (f) D10 25X coverage technical replicates. (g) Bars indicate correlation between replicates calculated for 65 replicate pairs from TKO library (23) and 3 from Brunello library (24) screens. Stars indicate pairwise correlations from (b) for FLI-seq (red) infection replicates or (blue) technical replicate libraries. (h) Scatter plot indicates frequency of sgRNAs with indicated read coverage for an example replicate pair from TKO library screening (23). This pair had median correlation across the 68 pairings considered in (g). For all, pearson correlation (R) indicated is calculated based only on sgRNAs with RPM>1 in at least one of the two datasets.

**Supplemental Figure 2.**
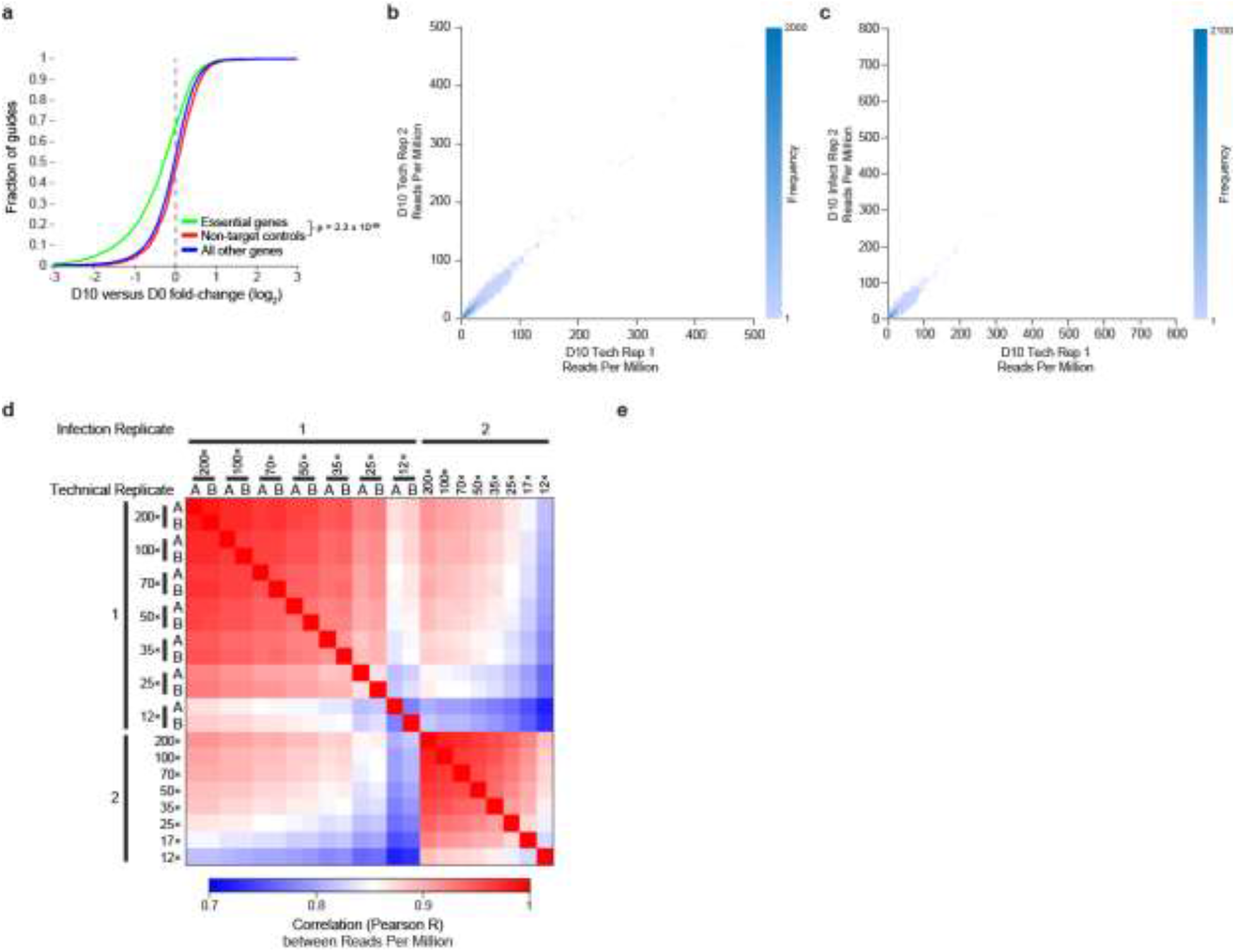
Reproducibility testing for improved library preparation for CRISPR/Cas9 pooled screens. (a) Cumulative distribution plots indicate fold-change between Day 0 and Day 10 observed for (green) annotated essential genes, (red) non-target controls, and (blue) all other genes. Significance was determined by Kolmogorov-Smirnov two-sided test. (b-c) Scatter plots indicate the density of sgRNAs showing the indicated normalized read coverage in (b) D10 technical replicates and (c) D10 infection replicates. See Fig. 2c-d for comparisons plotted on logarithmic scale. (b) Heatmap indicates correlation (Pearson R) between reads-per-million normalized sgRNA counts from technical or biological replicate samples as indicated, using only sgRNAs with RPM>1 in at least one dataset. See Fig. 2b for correlations calculated on log_2_-transformed normalized read density.

To confirm that the FLI-seq approach decreases variability compared to previous approaches, we compared its reproducibility to that observed in previously published CRISPR lethality screens. Considering 65 replicate pairs from TKO library (23) and 3 from Brunello library (24), we observed a median Pearson correlation of 0.65 (Fig. 2g-h). Indeed, the 90^th^ percentile correlation (R=0.76) was less than that observed with FLI-seq performed with final library readout of only 35x coverage (Fig. 2g). Thus, these results indicate that the FLI-seq method can yield highly reproducible library readouts from CRISPR/Cas9 pooled screens, with higher reproducibility and lower coverage than is frequently used in current procedures.

### Identification of effectors of DNA polymer micelle (DPANP) uptake

The ability of DPANPs to carry functional nucleic acids into the cytosol of cells and modulate mRNA expression of target genomes without transfection or other helper reagents makes them potential therapeutic nucleic acid carriers, but little is known about how they are able to access the cell interior (Fig. 3a). We previously showed that incorporation of a cyanine 5 (Cy5) dye into the DNA sequence at the end (3’) of a DPANP structure would enable visualization of micelle uptake into live cells (5). We identified regulators of DNA polymer micelle uptake using cellular Cy5 signal as a reporter for micelle uptake, as genetic or other cellular manipulations that decrease uptake will decrease the fluorescence readout (Fig. 3b). To enable this approach, we generated a modified DPANP micelle comprised of a 20-unit hydrophobic ROMP polymer covalently attached to a 30-nucleotide single stranded DNA sequence, with a cy5 label embedded via phosphodiester linkage within the strand backbone towards the 3’ end (Sup. Fig. 3a). The length of the sequence was chosen to be on par with an antisense strand, which is typically in the 20nt range, and is within facile synthesis range for solid phase phosphonamidite coupling procedures (which typically suffers in yield after 50 or more nts). The DNANP oligo sequence was chosen to be complementary to a sequence not expressed in mammalian cells (GFP), to minimize any potential binding to native sequences in the cell and includes the addition of T and A spacers at the polymer interface.

**Figure 3.**
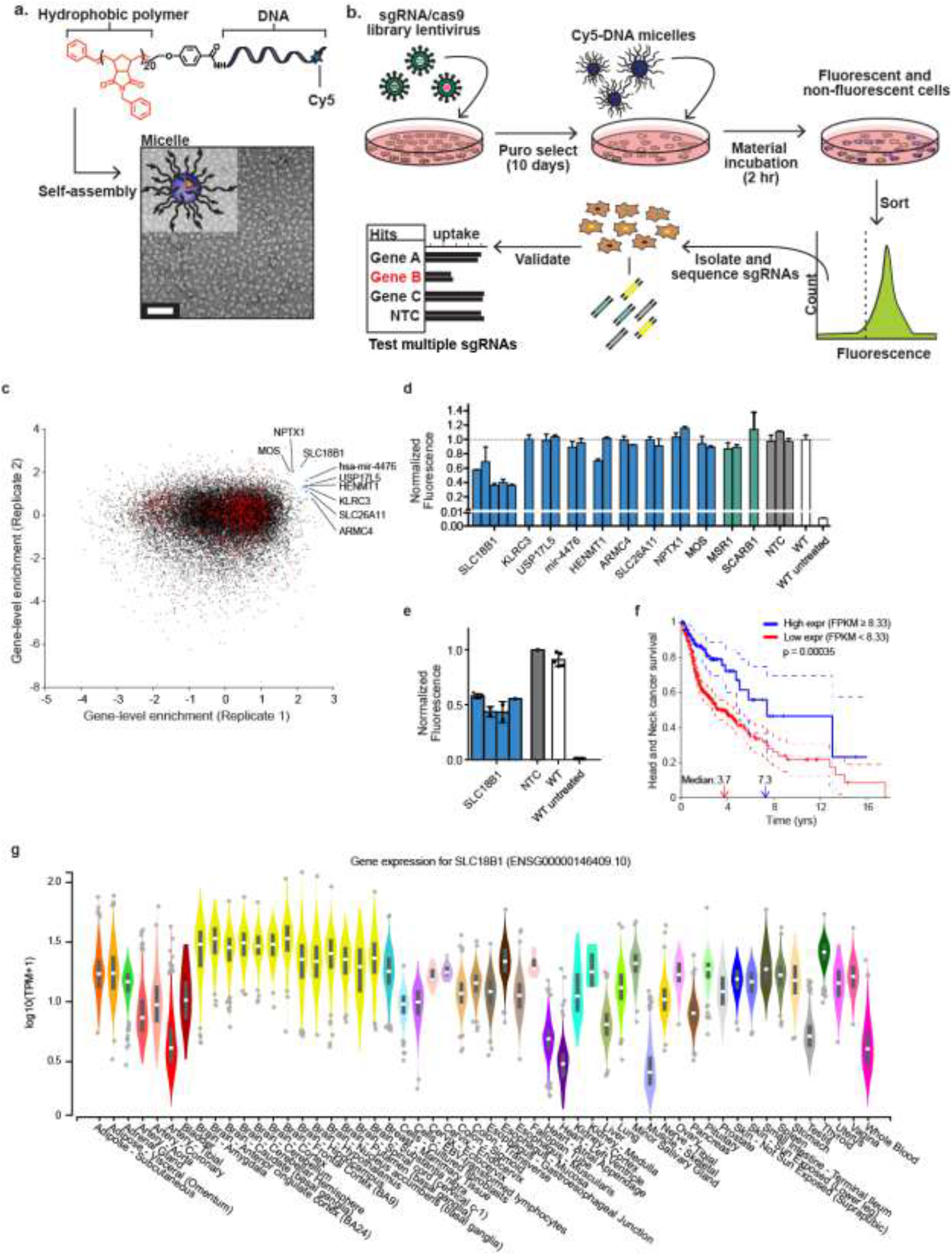
Screen for nanoparticle uptake identifies SLC18B1 as a novel effector. (a) Unimer and self-assembled structure of DPANPs. TEM of DPANPs shows discreet, ~20 nm micelles. Scale bar 100 nm. (b) Schematic of pooled screen. Whole-genome pooled sgRNA library lentivirus was transduced into HEK293T cells and viral infection was selected for 10 days with Puromycin. On Day 10, cells were treated with Cy5-labeled micelles, followed by FACS analysis to isolate the bottom ~2.5% fluorescent population as uptake-deficient. After library preparation, sgRNA enrichment in uptake-deficient versus unsorted population was used to identify candidate genes. (c) Scatter plot indicates gene-level sgRNA enrichment (z-score) from two biological replicate screens performed with the Gecko “A” library. Black points indicate genes, and red points indicate non-targeting controls. Nominally significant hits are indicated in blue. (d) Bars indicate fluorescent micelle uptake in single-knockout validation experiments. Each bar indicates an independent sgRNA sequence, and error bars indicate standard deviation from replicate measurements. (e) Bars indicate uptake observed in four clonal populations from SLC18B1 CRISPR/Cas9 knockout (with knockout confirmed by genomic PCR and sanger sequencing). (f) Kaplan-Meier plot shows (y-axis) fraction of patients surviving for (x-axis) indicated time, with patients separated by (blue) high and (red) low expression of SLC18B1 from TGCA data for head and neck cancer. Dotted lines indicate 95% confidence intervals, and censored datapoints are indicated by vertical lines. Plots were generated with KMplot package in MATLAB. (g) Violin plots indicate expression of SLC18B1 observed across 54 tissues profiled by the GTEx consortium, with box plot indicating median and 25^th^ to 75^th^ percentile.

**Supplemental Figure 3.**
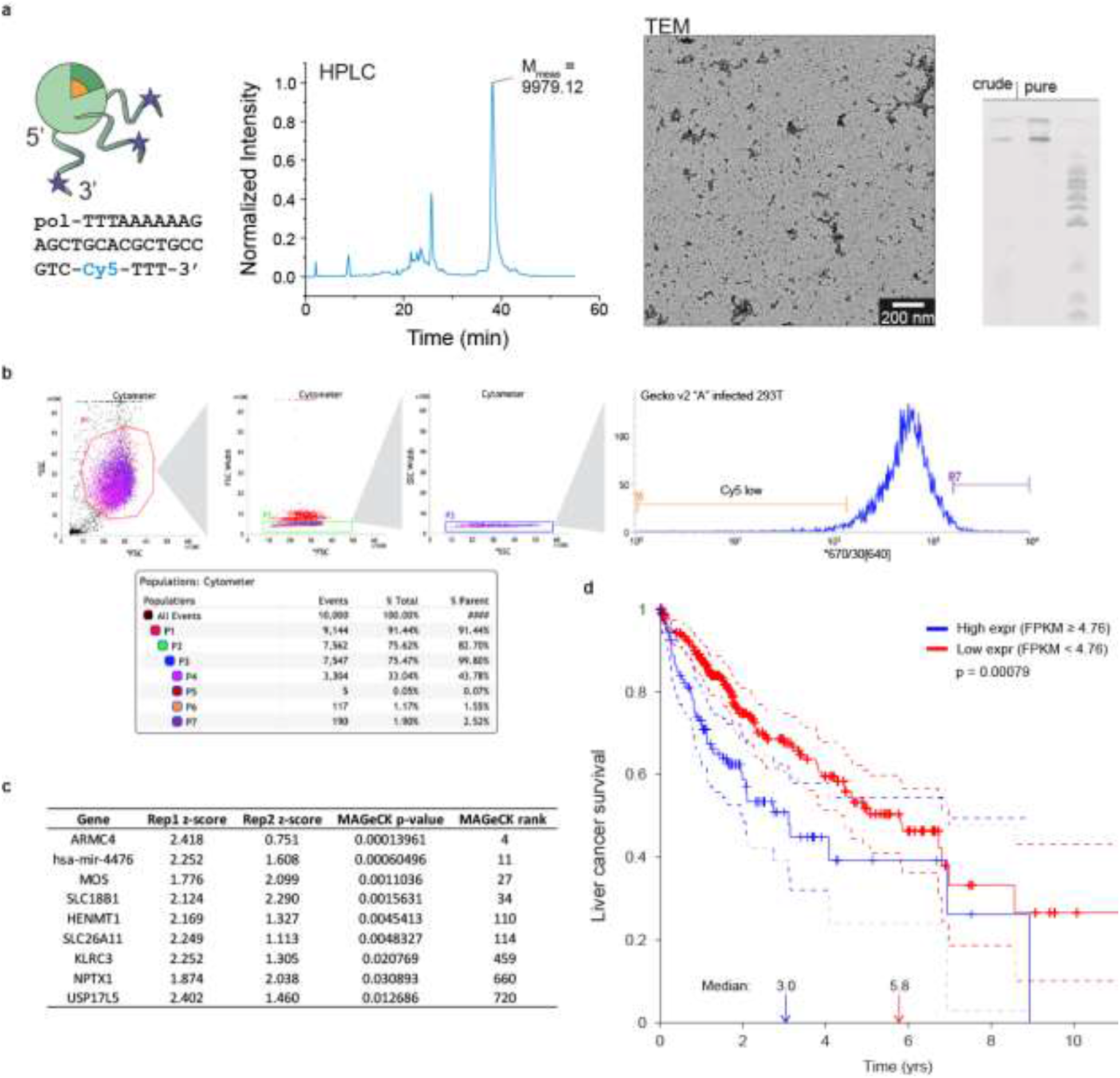
Screen for nanoparticle uptake identifies SLC18B1 as a novel effector. (a) From left to right: structure and sequence of DPANPs, pol = polymer. HPLC and corresponding mass (obtained via MALDI-TOF) of the Cy5 labeled DNA sequence. TEM micrograph of negatively stained DPANP micelles. PAGE gel of crude and purified DPANPs, visualized using ethidium bromide staining. (b) Fluorescence Activated Cell Sorting (FACS) gating strategy to select Cy5-low (P6; putative uptake-deficient) and Cy5-high (P7; putative uptake-increased) 293T cells that were infected with pooled CRISPR library. Data shown is an example quantitation from 10,000 sorted cells. (c) Ranking of selected genes in initial analysis (z-score relative to non-targeting controls) and re-analysis with MAGeCK comparing Cy5-low versus unsorted cells. (d) Kaplan-Meier plot shows (y-axis) fraction of patients surviving for (x-axis) indicated time, with patients separated by (blue) high and (red) low expression of SLC18B1 from TGCA data for liver cancer. Dotted lines indicate 95% confidence intervals, and censored datapoints are indicated by vertical lines. Plots were generated with KMplot package in MATLAB.

We performed two replicate infections of the Gecko ‘A’ library in 293T cells, and after 10-12 days of selection cells were treated with Cy5-labeled micelles followed by fluorescence activated cell sorting (FACS). As cells with the lowest Cy5 signal would contain sgRNA targeted to genes that mediate uptake of the nanomaterials, we selected the lowest 1-3% (Replicate 1; 694k out of 56M total input cells, Replicate 2; 1.71M out of 96.2M total input cells) as the ‘uptake deficient’ Cy5-low population (Sup. Fig. 3b). Genomic DNA was extracted from these populations as well as 7 million (Replicate 1) and 16 million (Replicate 2) unsorted cells as a control (100- and 240-fold library coverage respectively), and high-throughput libraries were generated using the FLI-seq method. We developed our own metric for identifying reproducibly enriched targets by calculating a z-score for the fold-enrichment observed for sgRNAs targeting each gene relative to 1000 included non-targeting sgRNAs (Fig. 3c, Supplemental Table 2, 3). Using this approach, we identified several potential candidates for effectors of micelle uptake, including both potential direct interactors (solute carrier proteins SLC18B1 and SLC26A11) and indirect effectors (miR-4476) (Fig. 3c). Repeating the analysis using the MAGeCK analysis pipeline (25) revealed generally similar enrichments, including identification of SLC18B1 as among the most significant candidates (Sup. Fig. 3c). We note that although the screen was designed to identify knockouts with near-complete loss of uptake, our later observation of more modest uptake deficiencies upon individual factor knockout may explain the overall low reproducibility observed in the genome-wide screen (Fig. 3c).

### SLC18B1 influences DNA polymer micelle uptake

We next prepared knockout cell populations for the top candidates to verify their respective influence on uptake. Two separate gRNAs were chosen per gene to prepare knockouts for validation: the most highly enriched guide from the uptake screen (GeCKO V2 ‘A’ library) and, when available, an independent guide from the more recently optimized Brunello human sgRNA library (26), which was not included in the initial screen. In cases where the Brunello guide was not available, the second most enriched guide from the screen was included. In addition to candidate hits from our screen, we prepared knockouts for two scavenger receptor proteins, MSR1 (rank 7976) and SCARB1 (rank 1904). These receptor proteins have been implicated previously in uptake of similarly structured nanoparticles by other methods (8, 27), but did not appear to have significant influence in our screen. After transduction and selection for infection, cells were incubated with micelles and fluorescence of each knockout was determined by flow cytometry. During this analysis, it became apparent that the highest ranked gene candidate, SLC18B1, displayed a significant (ca. 40-60%) reduction in fluorescence after material incubation. The other genes examined, as well as the genes implicated in the literature, showed no significant effect (Fig. 3d).

To further confirm the effect of SLC18B1, we tested an additional three guides (including guides not included in the original screen) which also showed a significant reduction in uptake compared to wildtype and non-targeting guide controls (Fig. 3d). Next, to account for potential heterogeneity of expression within the SLC18B1 knockout populations, we selected clones for four of the SLC18B1 guides and validated SLC18B1 mutation by targeted sequencing. Again, we observed upon micelle treatment that these cells demonstrated a reduction in fluorescence post-incubation consistent with the pooled SLC18B1 knockouts (Fig. 3e).

As one of the key advantages of DPANPs is to carry nucleic acids or other cargos into cells, the ability to differentially target cells based on SLC18B1 expression would present a potential avenue for future development of targeted therapies. To explore the potential for such approaches, we queried whether altered SLC18B1 expression was associated with disease progression in the Cancer Genome Atlas (TGCA) resource (28). Notably, we observed that higher expression of SLC18B1 served as a significant positive prognostic marker for head and neck cancer (Fig. 3f) and negative marker for liver cancer (Sup. Fig. 3c). Considering overall tissue-level expression from the GTEx consortium, SLC18B1 shows particularly higher expression across a variety of brain sub-regions, with variable expression in other tissues (Fig. 3g). Thus, not only does SLC18B1 represent a novel effector of DPANP uptake, the variable expression suggests that it may serve as an entry point into developing targeted DPANP therapies for different tissues or cell types.

## Discussion

The development of genome editing methods has revolutionized the ability to perform pooled forward genetic screens for a host of phenotypes. With an ever-increasing toolbox of reagents to perform gene knockouts, inhibitions, and over-expressions, pooled screening approaches are now within the capabilities of researchers across a host of fields (29). This has led to a greater need for improved and simplified methods to read out screen results and identify candidate hits in a robust and experimentally tractable manner. In particular, recent work has shown the power of combinatorial screens in which each cell is infected with dual sgRNAs to assay for genetic interactions (30), and saturation mutagenesis screens, in which a genomic region is tiled with sgRNAs to assay the presence and effect of functional elements (31). Both would require orders of magnitude deeper coverage than traditional loss-of-function screens to expand beyond small targeted panel approaches.

The nature of pooled screening requires amplifying a single ~200nt region per cell, leading to screens that require amplification from tens-to hundreds of micrograms of genomic DNA. Inhibitory effects of high DNA concentration per PCR have led to a variety of solutions, ranging from simply pooling hundreds of PCR reactions to utilizing restriction enzyme sites present in the lentiviral backbone constant regions flanking the sgRNA to perform DNA gel electrophoresis and size selection to remove undesired gDNA (19). However, these approaches can be both expensive and have significant handling challenges when scaled to large screens. To address this challenge, we developed a pulldown-based approach utilizing biotinylated RNA oligonucleotides complementary to the backbone constant regions flanking the sgRNA site. We observed that we could recover the majority of sgRNA-containing fragments while removing over 99% of gDNA, enabling a typical screen to be performed in a single PCR reaction. Using both technical (library preparation) and infection replicates, we show that the high recovery leads to high reproducibility, potentially enabling lower coverage screens.

With this preparation method for identifying modulators of a novel phenotype in hand, we turned to the challenge of determining effectors of nanomaterial uptake. Understanding interactions at the nanomaterial-cell level has been chronically challenging for researchers from both a synthetic and biological perspective. The current paradigm for understanding nanoparticle uptake typically involves employing pharmacological inhibitors to block general uptake pathways (32). However, these studies require prerequisite knowledge about likely uptake mechanisms, and the small molecule inhibitors often suffer from a lack of specificity, off-target effects and toxicity (33, 34). Furthermore, prior nanoparticle uptake studies using inhibitors suggest the chemical and geometric makeup of nanoparticles significantly alters their uptake profile, although how these features influence uptake remains unclear (35). Given the increasing abundance of nanoparticle-based therapies as the focus of basic research in nanomedicine broadly, or those entering clinical trials, a robust, modular screening approach to elucidate specific interactions between materials and their intended cellular targets is sorely needed (36, 37).

For the DNA-polymer amphiphile nanoparticle (DNANP) system, we identified Solute Carrier (SLC) factor SLC18B1 as a novel mediator of micelle uptake. SLC18B1 is one of over 400 solute carrier membrane transport proteins in the human genome, and despite numerous links to a variety of diseases only a small number of SLC proteins have been studied (38). SLC18B1/C6ORF192 was initially identified as a divergent member of the SLC16/17/18 family with expression across many tissue types but particular localization to secretory vesicles in the nervous system (39, 40). Although it has not previously been studied for its role in the uptake of nanoparticles, SLC18B1 has been characterized for the transport of polyamines, including spermine and spermidine (40). We hypothesize that the various primary and secondary amine groups displayed by the nucleotides at high density on the DPANP corona may mediate its interactions with the transporter.

The broad expression of SLC18B1 across many tissues, with particularly high expression throughout brain subregions, suggests that SLC18B1 may represent a mode of uptake in many tissues. However, if the 10-fold change in RNA expression of SLC18B1 is recapitulated in functional SLC18B1 protein on the membrane, it could suggest that SLC18B1-dependent micelle uptake will be highly tissue-dependent. It remains an open question whether this variability will be sufficient to drive the creation of tissue-targeted therapies. However, these data suggest the screening approach presented herein will be a powerful tool in identifying biomarkers for designing targeted molecules and materials.

The fluorescence-based screen selected for cells that were deficient in micelle uptake, with validation experiments with SLC18B1 repeatedly showing a 40-60% decrease in uptake. This suggests that there remain other SLC18B1-independent mechanisms of DPANP cellular entry. Although we were unable to validate other hits from the initial genome-wide screen, targeted screening of transporters (particularly in SLC18B1-deficient cell lines) may represent a focused approach to continue to explore the full set of modulators of DPANP micelle uptake. In addition, for these materials and other future materials or molecules analyzed via this approach, modifications to the particle surface could also serve to narrow in on which residues are important for transporter interactions, leading to structure function relationships at the bio-nanointerface. Such relationships typically remain difficult to probe, making the development of transporter targeted technologies from biomaterials to molecules difficult, where predictable performance guided by mechanistic understanding remains elusive.

## Methods

### Nanomaterial synthesis

#### Polymer synthesis

Reagents were purchased from commercial sources and used without further purification. The hydrophobic polymer portion of the DNA-polymer amphiphile was prepared via ROMP using a norbornene-phenyl monomer ((N-benzyl)-5-norbornene-exo-2,3-dicarboximide), carboxylic acid chain transfer agent (4,4’-(but-2-ene-1,4-diylbis(oxy))dibenzoic acid), and the ruthenium initiator [(IMesH_2_)(C_5_H_5_N)2(Cl)2Ru=CHPh], which were synthesized according to literature procedures (41–43). Monomer (500 – 600 mg) and initiator (1/20 molar ratio) were added to separate, dry Schlenk flasks with stir bars charged with N_2_ and dissolved in dry CDCl_3_ (3 mL and 0.5 mL, respectively). Once dissolved, the initiator solution was added to the monomer solution via cannulate with stirring. After 20 minutes, a 50 μL aliquot was removed and set aside for characterization by SEC-MALS. To the remaining reaction, 2 equivalents of chain transfer agent (with respect to initiator) in dry DMF was added and allowed to react for 45 minutes. The polymer was purified by precipitation three times in cold MeOH, then the pellet was redissolved in DCM and centrifuged to remove excess chain transfer agent. The supernatant was concentrated to dryness and run on a column (mobile phase 4% MeOH in DCM) and fractions containing the polymer product were combined and concentrated to dryness. Polymer dispersity and molecular weight were measured using a Phenomenex Phenogel 5μ 10, 10k-1000k, 300 × 7.80 mm (0.05 M LiBr in DMF)) using a Shimadzu LC-AT-VP pump equipped with a multiangle light scattering detector (DAWN-HELIOS: Wyatt Technology), a refractive index detector (Wyatt Optilab T-rEX), and a UV-vis detector (Shimadzu SPD-10AVP) normalized to a polystyrene standard. Polymer dispersity was measured to be 1.005; M_n_ = 5212 g/mol. Successful cross-metathesis with the chain transfer agent was monitored by ^1^H NMR spectroscopy on a Varian Mercury Plus spectrometer (400 MHz) by observing the shift of the alkylidene proton.

#### DNA synthesis

The hydrophilic DNA component of the amphiphile was synthesized using standard phosphoramidite coupling on an Applied Biosciences 394 automated synthesizer with a 1000 Å CPG column (Glen Research) on a 1 μmol scale. Phosphoramidite monomers (bz-dA-CE #10-1000-10, Ac-dC-CE #10-1015-10, dmf-dG-CE #10-1029-10, dT-CE #10-1030-10, Fluorescein-dT-CE #10-1056-90 and Cyanine 5 Phosphoramidite #10-5915-95) were purchased from Glen Research and dissolved in dry solvent according to manufacturer instructions. Synthesis conditions include activator 4,5-Dicyanoimidazole (Glen Reasearch), 0.02 M iodine in THF/pyridine/water (Glen Research) as the oxidizer, 3% Trichloroacetic acid (Glen Research) as the deblocking solution, and capping mixtures THF/Pyridine/Ac_2_O (Glen Research) and 16% 1-Methylimidazole in THF (Glen Research). A 5’-amino modifier (Glen Research) was coupled onto the end of sequences for subsequent conjugation with the polymer. Aliquots of each sequence (2-5 mg) were cleaved from the solid support and deprotected (concentrated ammonia for 24-36 hr) for analysis. The MMT protecting group of the amino modifier was left intact to act as a drag-tag for HPLC purification. HPLC purified sequences were desalted using C18 resin (Ziptips, Millipore) and confirmed by MALDI-TOF using 2’,4’,6’-Trihydroxyacetophenone/ammonium citrate and 3-hydroxypicolinic acid as a matrix.

#### Micelle formation

DNA was then conjugated to polymer on solid support. Polymer (30 mg) was dissolved in 100 μL dry DMF with 5 μL DIPEA. Coupling agent (HATU, 0.9 equivalents with respect to polymer) was added to the solution and allowed to activate for 10 minutes. Concurrently, the MMT protecting group on the 5’-amino terminus of the DNA was deprotected using trichloroacetic acid followed by rinsing and drying the DNA on solid support with DCM and Ar, respectively. The DNA on CPG beads were then added to the activated polymer solution in a microcentrifuge tube and allowed to react at room temperature on a shaker. After two hours, the reaction was centrifuged to pool the CPGs with DNA-polymer conjugates and the supernatant was removed. CPGs were washed three times with 1 mL DMF by centrifugation, then a freshly activated solution of polymer (prepared in identical fashion as above) was added to the washed beads for a second reaction, which was left on a shaker overnight. The conjugated CPG was then washed in the same fashion as described previously, and on the last wash the beads were returned to the manufacturer CPG column and washed by flowing 15 mL DMF, followed by 15 mL DCM through the column to remove any unreacted polymer. After drying under Ar, the CPG beads were transferred to 2 mL plastic vials and cleaved/deprotected with 0.5 mL concentrated ammonia for 24-36 hours. Micelles from the DNA-polymer conjugate are spontaneously formed in the aqueous deprotection solution, and after removing the beads by filtration and washing with H_2_O, DMSO, and formamide, the filtrate containing micelles and any unreacted DNA was dialyzed into nanopure H_2_O using 10k MWCO snakeskin dialysis tubing (Thermo Scientific). Crude micelle solution was then concentrated to 1 mL under reduced pressure, and injected on SEC-FPLC (HiPrep 26/60 Sephacryl S-200 High Resolution packed column with mobile phase of 50 mM Tris pH 8.5 on an Akta purifier (Pharmacia Biotech) running a P-900 pump and a UV-900 UV-Vis multi-wavelength detector) to purify. Pure micelles were then dialyzed into nanopure H_2_O and concentrated under reduced pressure. Characterization of micelles was performed by denaturing PAGE (15% separating, 7% stacking, 1×TBE, 8 M Urea) to confirm purity, and TEM to confirm morphology. TEM samples were prepared by glow-discharging grids (formvar/carbon-coated, 400 mesh copper, Ted Pella) at 20 mA for 90 seconds using a s4 Emitech K350 glow discharge unit followed by treatment with 250 mM CaCl_2_. After blotting away excess salt solution, 3.5 μL micelle sample was added to the grid and allowed to dry for 10 minutes and then washed by passing 3 drops of glass distilled H_2_O over the grid followed by 3 drops of 1% w/w uranyl acetate stain, the excess of which was immediately blotted away using filter paper. The sample-loaded grid was then loaded onto the microscope for imaging.

#### Cell culture

HEK293T cells were cultured at 37°C and 5% CO_2_ in complete medium (Dulbecco’s Modified Eagle Medium, Life Technologies) supplemented with 10% fetal bovine serum (OmegaScientific), and 1X penicillin/streptomycin (Corning). Cells were maintained in 10 cm petri dishes and passaged at ~75 – 90% confluency every 3 – 5 days.

#### Amplification of pooled sgRNA plasmid DNA

Human GeCKOv2 CRISPR knockout pooled library was a gift from Feng Zhang (Addgene #1000000048) (16). To amplify the A sub-pool, 400 ng of pooled plasmid DNA was electroporated into Endura (Lucigen) E. coli (performed as 4 electroporations of 2 ul (100 ng) pooled plasmid DNA into 25 uL cells), 2 mL of recovery buffer (Lucigen) was added, and cells were recovered at 37°C for 1 hour. Cells were then pooled, plated onto 245mm bioassay plates, and incubated at 33°C for ~16 hours. Cells were then scraped and maxiprepped (Qiagen) with a maximum of 450 mg bacteria per column. This same procedure was repeated for the B sub-pool. Colony counting estimated total recovery of 1.44×10^8^ and 9.6×10^7^ colonies for the A and B libraries respectively.

#### Preparation and titer of pooled library lentivirus

To prepare pooled library lentivirus, 6×15 cm plates of HEK293XT cells were grown to 40% confluency and transitioned to Opti-MEM reduced serum media for one hour. For each plate, transfection mix was prepared as one mixture of 2.5 mL Opti-MEM, 124 μL Lipofectamine PLUS reagent (ThermoFisher), 14.4 μg Gecko v2 plasmid DNA, 6.2 μg pMD2.G, and 9.3 μg psPAX2, and a second mixture of 2.5 mL Opti-MEM with 62.2 μL Lipofectamine 2000. The two mixtures were incubated for 5 minutes at room temperature, mixed, incubated for 10 minutes at room temperature, and then the 5 mL mixture was added dropwise to the 15 cm plate. Six plates each were transfected for both the A and B library sub-pools. After 6 hours, media was replaced with standard growth media (DMEM with 10% FBS).

Lentivirus was harvested 56 hours post-transfection by filtering media through 0.45 μm PES filter. Filtered media was then ultracentrifuged at 20,000 rpm for 2 hours at 4°C, and 100× concentrated lentivirus was resuspended in DMEM with 10% FBS, incubated overnight at 4°C, and aliquoted and stored at −80°C.

To test lentiviral titer, 1.5×10^6^ HEK293XT cells were plated per well of a 24 well plate in DMEM with 10% FBS supplemented with 8 ug/mL protamine sulfate. After 30 minutes, varying amounts of pooled library lentivirus was added and spinfection was performed by centrifuging plates at 37°C for 2 hours at 2,000 rpm, after which media was changed. The following day, each well was split into paired wells with and without puromycin. After 48 hours of selection, cell survival was determined by hemocytometer count, and viral amounts equivalent to multiplicity of 0.3 were identified and used for following experiments.

#### Pooled CRISPR/Cas9 screening

For full screen experiments, 24 wells of 3×10^6^ HEK293T were plated in 12 well plates in DMEM supplemented with 10% FBS and 8 μg/mL protamine sulfate. After 30 minute incubation, 3 μL lentivirus (equivalent to 0.3 MOI) was diluted in 100 μL of Opti-MEM and added dropwise to each well. Plates were then centrifuged at 2,000 rpm at 37°C for 2 hours, after which media was changed to standard DMEM supplemented with 10% FBS. 30 hours post-transduction, cells were passaged into media containing 1 μg/mL Puromycin. Cells were passaged for 10 days maintaining at least 65×10^6^ cells (1000× coverage) per passage. For library preparation experiments, cell pellets were frozen for gDNA extraction and library preparation at the indicated timepoints.

#### Micelle uptake CRISPR/Cas9 screen

For uptake screening, at 10 days post transduction, cells were treated with 0.1 nM (replicate 1) or 0.2 nM (replicate 2) Cy5-labeled micelles and incubated for 2 hours. To identify uptake-deficient cells, Fluorescent Activated Cell Sorting (FACS) was performed at the UCSD Human Embryonic Stem Cell Core Facility on a FACSAria2 sorter. Gates were set on the 640 laser (670/30 filter) to isolate the ~1% (~6.9×10^5^ cells out of 56 million sorted; replicate 1) and ~3% (1.7×10^6^ cells out of 96.2 million sorted; replicate 2) of cells with the lowest Cy5 signal (see Supplemental Figure 3a for example). Final sequenced libraries were prepared from a subset of these samples (Replicate 1: 1.8 μg gDNA equivalent to ~2.7×10^5^ cells; Replicate 2: 12 μg gDNA equivalent to 1.8×10^6^ cells). As an input, a population of untreated and unsorted cells was selected (Replicate 1: 7.1×10^6^ cells; Replicate 2: 16.1×10^6^ cells).

### FLI-seq library preparation

#### Probe generation

To generate antisense probes for targeted enrichment, two templates with T7 promoter sequences were generated by performing PCR amplification (Q5 Polymerase, NEB) off of GeCKO v2.0 plasmid DNA using H1T7 and H1R primers for probe 1 and primers T2RT7 and T2F for probe 2 (Supplemental Table 4). Probe 1 hybridizes to the negative strand of upstream sequence, whereas probe 2 hybridizes to the forward strand of the downstream sequence (Fig. 1b). PCR products were cleaned using AMPure XP beads (Beckman Coulter) and manufacturer protocol. In later experiments, amplified PCR products were re-amplified to generate greater probe amounts. Purified dsDNA templates were used as templates for T7 transcription reactions using HiScribe™ T7 High Yield RNA Synthesis Kit (NEB) and bio-16-CTP (Trilink) and bio-16-UTP (Roche) nucleotides at 1:10 ratio to normal nucleotides. After T7 transcription, RNA probes were purified using RNA Clean & Concentrator-25 kit (Zymo Research).

#### Enrichment for sgRNA genomic sequences and library prep

gDNA was isolated using the DNeasy Blood and Tissue Kit (Qiagen), including RNAse A treatment. gDNA was quantified by TapeStation 2200 (Agilent), and then fragmented by sonication (either Q800R2 Sonicator (Qsonica) or Bioruptor Plus (Diagenode)) to 800-1000bp fragments. After gDNA was denatured, 10% of biotinylated probes (by mass; e.g. 10 μg probes for 100 μg gDNA) were pre-coupled to Streptavidin beads (ThermoFisher) using the manufacturer recommended protocol, and were then hybridized to vector-containing gDNA fragments in 1× LiCl/Urea Buffer (the final 1× Buffer contains 25 mM Tris pH 7.4, 5 mM EDTA, 400 mM LiCl, 0.1% NP40, 0.1% SDS, 0.1% sodium deoxycholate, 1M urea) for 3 hours at 60°C. After hybridization, beads were rinsed with 1× LiCl/Urea Buffer at 45°C for 2-3 minutes. sgRNA-containing gDNA fragments were eluted by digesting RNA probes with Ambion^®^ RNase Cocktail™ (ThermoFisher) and by degrading RNA with NaOH (Sigma). gDNA fragments were purified using the DNA Clean & Concentrator-5 kit (Zymo Research).

Purified gDNA fragments were then PCR amplified (NEBNext Ultra II Q5, NEB) with FLI1F and FLIR primers in PCR1 (denaturation at 98°C for 30 seconds, followed by 6-9 cycles of denaturation at 98^°^C for 12 seconds, annealing at 69°C for 60 seconds, and extension at 72°C for 30 seconds, followed by a final elongation at 72°C for 60 seconds). PCR1 products were purified using AMPure XP beads at 1.6× ratio by volume and manufacturer protocol. After purification, PCR2 was performed with NEBNext Ultra II Q5 (NEB) using standard Illumina indexed primers (e.g. D501 and D701) as follows: denaturation at 98°C for 30 seconds, followed by 7-8 cycles of denaturation at 98°C for 10 seconds and annealing and elongation at 72°C for 40 seconds, followed by a final elongation at 72°C for 60 seconds). PCR2 products were purified using AMPure XP beads at 1.3× ratio by volume and manufacturer protocol. Libraries were quantified by Tapestation 2200 (Agilent) and sequenced on the HiSeq 2500 or 4000 platforms (Illumina). As these libraries contain identical sequences upstream of the variable sgRNA region, pools submitted for sequencing contained at most 20% FLI-seq libraries, with the remainder of the pool containing high-diversity samples (e.g. standard RNA-seq).

#### Analysis of pooled CRISPR/Cas9 screens

Sequencing reads were first processed by requiring the presence of flanking upstream (GTGGAAAGGACGAAACACCG) and downstream (GTTTT) sequences flanking a 20nt region. These 20nt regions were tallied, and then queried against sgRNA sequences present in the Gecko v2 “A” or “B” libraries. Per-sgRNA Reads Per Million (RPM) were calculated by adding a pseudocount of 1 read and normalizing against the sum of all sequenced reads that contained an exact sequence match to a sgRNA sequence in the queried library. Pair-wise correlations in log_2_(RPM) were calculated using only sgRNAs with RPM>1 in at least one of the two datasets.

For identification of differentially enriched sgRNAs upon micelle treatment, a pseudocount of one read added to both values and then fold-change was calculated between RPM values in Cy5-low sorted population relative to unsorted. Next, for each sgRNA a z-score was calculated by comparing the fold-change against the distribution of fold-changes observed for 1000 non-targeting sgRNAs. A gene-level z-score was then calculated using the Stouffer’s Z-score method. As an alternative approach, sgRNA counts (as described above) were used as input to the MAGeCK (v0.5.9.3) analysis tool (25). To validate depletion of essential genes, the CEG2 list of 684 core essential genes was obtained from Hart *et al*. (22).

To compare against previously published CRISPR/Cas9 screen reproducibility, sgRNA read counts were obtained for the TKO library from http://tko.ccbr.utoronto.ca, including 65 total pair-wise replicate comparisons (3 comparisons in DLD1 cells, 3 timepoints in GBM cells, 2 libraries with 5 timepoints in HCT116 (experiment 1), 2 libraries with 4 timepoints and 3 comparisons per timepoint in HCT116 (experiment 2), 2 libraries with 4 timepoints and 3 comparisons per timepoint in HeLa (excluding T18 for library 1, which was discarded due to abnormal correlation with T15), and 4 timepoints in RPE1 cells (23). Additionally, sequencing data for screening performed with the Brunello library in A375 cells was obtained (24). Processing was performed identically to described above by identifying reads containing flanking upstream (CGAAACACCG) and downstream (GTTT) sequences and zero mismatch alignment to an sgRNA in the Brunello library. Final read density was calculated by adding a pseudocount of 1 read and normalizing to reads per million based on the total number of zero-mismatch reads in the sample. Pair-wise correlations in log_2_(RPM) were calculated using only sgRNAs with RPM>1 in at least one of the two datasets.

### Validation of modulation of uptake by SLC18B1

#### Plasmids

LentiCRISPR v2 was a gift from Feng Zhang (Addgene plasmid #52961; http://n2t.net/addgene:52961; RRID:Addgene_52961). LentiCRISPR v2 plasmids were prepared and amplified according to established procedure (16). Sequences for the gRNAs were chosen based on: 1) highest enrichment in the pooled screen, or 2) an optimized sequence from the Brunello library (26). Five non-targeting control guides were also prepared as controls. Insert oligos were purchased from Integrated DNA Technologies (IDT). Post-amplification plasmid sequence was confirmed by Sanger sequencing (Genewiz) of the insert DNA using the LKO.1 5’ primer.

#### Virus production

HEK293Ts were seeded in 24-well format at 100k cells/well one day prior to transfection. Immediately before transfection, medium was removed and replaced with antibiotic-free medium. Packaging plasmid (psPAX2), envelope plasmid (pMD2.G), and transfer plasmid (LentiCRISPRv2) were mixed with Lipofectamine^®^ 2000 (ThermoFisher) in OptiMEM (ThermoFisher) according to the manufacturer protocol and incubated for five minutes prior to adding to wells. A total of 600 ng DNA was transfected per well, with a plasmid ratio of 4:3:1 (wt/wt/wt) packaging:envelope:transfer in a total volume of 50 μL OptiMEM. Virus was harvested at 48 hr and 72 hr post-transfection and filtered through 0.45 μm PVDF syringe filter units (Millipore).

#### Transduction

HEK293Ts were seeded in 24-well format at 100k cells/well one day prior to transduction. Immediately before transduction, medium was removed and replaced with antibiotic-free medium. To each well, 200 μL freshly filtered virus was added. The same transduction procedure was repeated with freshly harvested and filtered virus the following day. At ~ 95% confluency, transduced cells were seeded into 6-well format in 1.5 μg/mL puromycin (Life Technologies) containing medium. Cells were allowed to select in puromycin for 9 days, replacing medium or passaging every 3 – 4 days.

#### Cloning SLC18B1 knockouts

From the validation knockout populations generated in the previous section, cells were harvested and seeded in 10 cm petri dishes at a density of 800 – 1000 cells/plate. After 3 – 5 days, five colonies from each knockout were selected and gently removed from the bottom of the dish using a 200 μL pipet and transferred to individual wells of a 96-well plate. At ~ 80% confluency, the cells were passaged. At the next passage, cells were transferred to 48-well plates. mRNA was extracted from the cells and treated with DNaseI (NEB) followed by reverse transcription with Superscript III (Invitrogen) using dT(20) primers according the manufacturer’s instructions. The abundance of target cDNA relative to the housekeeping control GAPDH was measured using qPCR. Cells that demonstrated successful and complete knockout of the target gene were maintained in culture and used in material uptake studies.

#### Uptake

Transduced HEK293Ts were seeded in 96-well format at 30k cells/well one day prior to treatment with micelles. Medium was replaced with 50 μL micelle-containing OptiMEM at a concentration of 0.1 nM (or 0.0 nM for control wells) with respect to micelle (assuming ~ 200 unimers per particle) (11). Treated cells were incubated for 2 hr at 37 °C, then the OptiMEM was removed and cells were washed with DPBS and trypsonized. Cells were resuspended in100 μL ice cold DPBS and immediately analyzed by FACS. FACS was performed on a BD FACSCanto and laser settings remained consistent between all wells/runs.

## Acknowledgements

The results shown here are in part based upon data generated by the TCGA Research Network: https://www.cancer.gov/tcga. The Genotype-Tissue Expression (GTEx) Project was supported by the Common Fund of the Office of the Director of the National Institutes of Health, and by NCI, NHGRI, NHLBI, NIDA, NIMH, and NINDS. The data used for the analyses described in this manuscript were obtained from the GTEx Portal on 9/29/19. ELVN was a Merck Fellow of the Damon Runyon Cancer Research Foundation (DRG-2172-13) and was supported by the NHGRI (K99HG009530). The work is partially supported by NIH grants HG004659 and NS103172 to GWY. In addition, the authors thank the NSF for support of this research through (DMR-1710105).

## Data Availability

CRISPR screen sequencing data has been deposited at the Gene Expression Omnibus (accession GSE157524).

## Code Availability

Scripts for analysis of CRISPR screen high-throughput sequencing data are available upon request.

